# HOPMA: Boosting protein functional dynamics with colored contact maps

**DOI:** 10.1101/2020.12.31.424963

**Authors:** Elodie Laine, Sergei Grudinin

## Abstract

In light of the recent very rapid progress in protein structure prediction, accessing the multitude of functional protein states is becoming more central than ever before. Indeed, proteins are flexible macromolecules, and they often perform their function by switching between different conformations. However, high-resolution experimental techniques such as X-ray crystallography and cryogenic electron microscopy can catch relatively few protein functional states. Many others are only accessible under physiological conditions in solution. Therefore, there is a pressing need to fill this gap with computational approaches.

We present HOPMA, a novel method to predict protein functional states and transitions using a modified elastic network model. The method exploits patterns in a protein contact map, taking its 3D structure as input, and excludes some disconnected patches from the elastic network. Combined with nonlinear normal mode analysis, this strategy boosts the protein conformational space exploration, especially when the input structure is highly constrained, as we demonstrate on a set of more than 400 transitions. Our results let us envision the discovery of new functional conformations, which were unreachable previously, starting from the experimentally known protein structures.

The method is computationally efficient and available at https://github.com/elolaine/HOPMA and https://team.inria.fr/nano-d/software/nolb-normal-modes.

## 1. Introduction

Proteins and their complexes are intrinsically flexible, and this flexibility is tightly linked to their biological functions (1, 2). The shape a protein adopts in solution and the way it moves is governed by a myriad of interatomic forces within the protein and with the surrounding solvent. Despite this high complexity, many motions relevant to protein functioning can be fairly approximated by a few low-frequency modes computed using the Normal Mode Analysis (NMA) and characteristic of the protein’s geometrical shape (3–7). This observation has led to the development of many NMA-based methods to predict crystallographic temperature factors (8–12), refine crystallographic structures (13, 14), interpret or fit atomistic structures into cryo-EM maps (15–24) or one-dimensional scattering profiles (25–27), and predict the structures of protein complexes in combination with docking (1, 28–38), among other applications.

While the NMA formalism may be used with standard semi-empirical potential functions, highly simplified potentials, called *Elastic Network Models*, where neighboring atoms are connected by harmonic springs yield a good approximation of the low-frequency normal modes and are thus commonly used (3–5). They have proven useful in modeling protein transitions (39–45), sampling protein conformational space (34, 46–49), partitioning proteins into rigid bodies (50–52), scanning single and double mutations (53–57), modeling allosteric signal propagation (58–60), and more. They have also helped to characterize the relationship between protein structural dynamics and the evolution of protein sequences (61–66).

The suitability of the NMA to model conformational dynamics crucially depends on the type of motions involved and on the protein 3D shape used to compute the modes (67). Typically, less constrained conformations encode more information about the dynamical potential of the protein than more constrained conformations. This is particularly visible when predicting a conformational transition from a closed to an open state. Indeed, opening the protein is significantly more difficult than closing it (3, 68). Lowering the distance cutoff used to determine whether two atoms are connected in the network may help to partially solve the problem. However, there is no universal optimum value applicable to any protein, and even within a given protein, it may be far from trivial to set a value that alleviates the constraints in some regions while avoiding sub-critical connectivity in others.

Several works have proposed to modulate the spring’s stiffness depending on the type of interatomic connections or the local packing density (10, 69, 70). The resulting hybrid or multi-scale networks yield more accurate predictions of the temperature factors, but this gain is at the expense of a loss of universality and simplicity. Complications may arise, for example, from asymmetries in the definition of the network (70). Another strategy consists in updating the elastic network iteratively, either by re-building a network from a conformation obtained by deforming the initial network (68, 71), or by modifying the equilibrium distances on-the-fly, for example to fit the network into a low-resolution map (18, 72).

Here, we report on a method enhancing protein elastic networks dynamical potential by exploiting patterns in protein contact maps. We build smoothed binarized colored contact maps from inter-residue distances and identify the patches in these maps that correspond to small contact areas between protein segments. We remove these patches and compute negative images of the filtered contact maps defining pairs of protein regions that should not be linked by any spring. Contrary to protein domain detection methods (73–76), we do not attempt to partition the protein in a set of rigid bodies but focus on detecting the interatomic connections that lock the protein and hinder its motions. Our approach, called HOPMA (proteiN cOloRed MAps), is conceptually simple, has only a few parameters and relies solely on the 3D information of the input structure. We show that HOPMA facilitates protein conformational space sampling by computing more than 400 protein functional transitions. We systematically assess HOPMA’s contribution to the description of the transitions by comparing the results obtained when starting from a modified protein elastic network, where the contacts identified by HOPMA have been removed, with those obtained from “classical” elastic networks, only defined based on a distance cutoff.

There are multiple ways to compute protein transitions with NMA, for example, using normal modes in internal coordinates (77–81). Here, we used the NOnLinear rigid Block (NOLB) NMA, a Cartesian nonlinear approach that extrapolates the instantaneous eigenmotions as a series of twists (82). NOLB is computationally very efficient, as it can be applied to systems as large as viral capsids or ribosomes at all-atom resolution on a personal laptop (82). It produces conformations with high stereochemical quality and is able to predict motions with different degrees of collectivity, from localized and disruptive motions to highly collectives ones (68). We place ourself in the context where we know the target structure and we use this knowledge to determine the amplitudes of the initial structure’s normal modes. This set up allows obtaining the optimal (or close-to-optimal) transitions within our framework. In practical applications, when the target is not know, the sampling may be guided by additional information (83, 84), such as experimental small-angle scattering profiles (25–27, 85, 86), low-resolution electron density maps (15–22), intensities from crystallographic scattering experiments (87, 88), complementarity of the molecular shapes upon binding (28–36), or other type of data such as contacts inferred from coevolutionary signals extracted from protein sequences (89–91).

## 2. Materials and Methods

### Datasets

We considered four reference protein benchmark sets, namely the iMod benchmark (92), the HR-PDNA187 benchmark (93), the Protein-Protein Docking Benchmark v5 (PPDBv5) (94), and the dataset reported in **(author?)** (90) (**Table 1**). We used the iMod benchmark as the validation set and the other benchmarks as testing sets.

**Table 1.** Benchmark sets

|  | #(structures) | #(transitions) | RMSD range (in Å) | types of transitions |
| --- | --- | --- | --- | --- |
| iMod (92) | 23 | 46 | 1.28 - 13.55 | opening/closing |
| HR-PDNA187 (93) | 11 | 11 | 2.22 - 28.37 | DNA binding |
| PPDBv5 (94) | 68 | 68 | 2.01 - 32.79 | protein binding |
| AlterBench (90) | 224 | 282 | 3.85 - 34.49 | diverse |

### Opening/closing set

The iMod benchmark, available at http://chaconlab.org/multiscale-simulations/imod/imod-donwload/item/imod-benchmark, was previously used to assess coarsegrained elastic network model-based flexible fitting methods (95). It comprises 23 proteins, each given in “open” and” closed” conformations, and represents a wide variety of macromolecular motions, predominantly hinge motions but also shear and other complex motions. The structures come from the molecular motions database MolMovDB (96). All of them have less than 3% Ramachandran outliers (as computed by the MolProbity program (97)), do not have any broken chain or missing atom.

### Unbound-to-bound sets

The HR-PDNA187 benchmark, available at http://www.lcqb.upmc.fr/PDNAbenchmarks/, was recently compiled to assess methods predicting DNA-binding patches at the surface of proteins. It comprises 187 high-quality structures (better than 2.5 Å resolution) of protein-DNA complexes and covers all major groups of protein-DNA interactions. The proteins have less than 25% sequence identity. 82 complexes come along with the unbound forms of the proteins. We extracted 11 monomeric unbound-bound (free-complexed) pairs with C_*α*_ RMS deviations between the two states above 2 Å.

The PPDBv5 benchmark, available at https://zlab.umassmed.edu/benchmark/, is well suited for assessing the range of applicability of flexible docking methods (98). It contains 230 protein complexes with at least one of the partners solved in both bound (complexed) and unbound (free) states. All structures have a resolution better than 3.25 Å, and some of them contain more than one chain. We extracted 68 singlechain proteins with C_*α*_ root-mean-square deviation (RMSD) between the two states above 2 Å.

### Large diverse set

The dataset reported in **(author?)** (90) was originally compiled to assess the ability of coevolutionary signals to guide the discovery of alternative functional protein conformations. It comprises 92 proteins sharing less than 70% sequence identity and corresponding to 224 protein chain structures. The motions exhibited by these structures are large (at least 3 Å RMSD) and of different types, namely open-close, rotation-close, rotation, concerted and other. The structures are grouped in 143 unique pairs, among which three are also found in the iMod benchmark (1akeA-4akeA, 1gggA-1wdnA, 1lstA-2laoA). For each pair, we considered both the forward and backward transitions, leading to a total of 286 transitions. We excluded four transitions because their initial structures (2ceaA, 3g7sB, 3rrmA and 1qmpB) displayed missing residues in the region serving as a hinge for the motion. Hence, the two parts of the protein moving with respect to each other were not connected by the protein backbone. We refer to this set as AlterBench.

### The proteiN cOloRed MAps (HOPMA) algorithm

HOPMA takes as input a protein 3D structure and gives as output a set of protein region pairs defining links that should be excluded in the elastic network model representing the input structure.

### From distances to contact propensities

Given a protein 3D structure, we first compute all pairwise C*α*-C*α* distances between the protein residues. Then, we convert these distances into contact propensities ranging between 0 and 1. The contact propensity *p*_*ij*_ between residues *i* and *j* is expressed as,

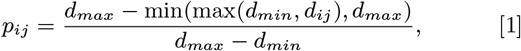

where *d*_*ij*_ is the distance between the C*α* atoms of residues *i* and *j*, and *d*_*min*_ and *d*_*max*_ are predefined minimal and maximal distance bounds. Intuitively, a pair of residue distant by less than *d*_*min*_ will be considered as very close to each other and their contact propensity will be equal to 1. By contrast, a pair of residues distant by more than *d*_*max*_ will be considered as very far from each other and their contact propensity will be null. Different cutoff distances, typically in the interval of 6–12 Å, have been proposed in the literature to define C*α*-C*α* contacts. Here, we set *d*_*min*_ = 3 Å and *d*_*max*_ = 15 Å.

### Detecting contiguous interaction patches

Next, we proceed through a smoothing of the contact propensity matrix. Specifically, the smoothed value 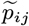 for the residue pair (*i,j*) is computed as the averaged contact propensity between the residues and their neighbours in the protein sequence:

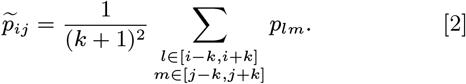

We empirically set the size of the smoothing window to *k* = 2, such that the average contact propensities are computed over 25 cells of the contact propensity matrix *P*. In practice, as the matrix *P* is symmetrical, the smoothed values are computed only for the cells located in its lower triangle.

Finally, we convert the smoothed real-value matrix 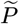 into a binarised matrix *B*, depending on a cutoff value *p*_*cut*_,

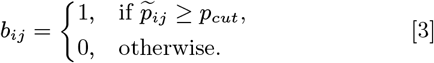

In this binary matrix, the cells with a value of 1 are considered as active and the others as inactive. We further colour the matrix by detecting contiguous patches of active cells and labelling each active cell with a unique patch identifier using a fast-scanning version of the Hoshen–Kopelman algorithm, also called the connected component labelling 2-pass algorithm (99, 100).

### Defining the set of excluded contacts

For each input structure, we compute two coloured maps with two different *p*_*cut*_ values, namely 0.001 and 0.1. The first value will lead to activating an (*i,j*) cell (*i*.*e*. setting its value to 1) whenever at least one of the residue pairs in the associated 25-cell window *w* = {(*l, k*) | *i* −*k* ≤ *l* ≤ *i* + *k, j* −*k* ≤ *m* ≤ *j* + *k*} is distant by less than 15 Å. The second value defines a more stringent activation criterion, which will be met for example when 3 residue pairs in the window are very close to each other (C*α* atoms distant by less than 5 Å). Within each coloured map, we select the contiguous patches smaller than (*k* + 1)^2^ cells, where *k* is the size of the smoothing window. Here, as *k* = 2, we select patches up to 625 cells, which typically corresponds to an interaction between two supersecondary structures. We filter out the selected patches from the binary contact matrix *B* by inactivating the corresponding cells (*i*.*e*. setting their value to zero). We compute the final matrix as the average of the two filtered matrices 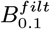 and 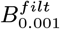 and we extract the 𝒮 = {(*s*_1_, *s*_2_), (*s*_3_, *s*_4_), …} of protein segment pairs defining the inactive regions (values lower than 1) of that matrix.

### Computing non-linear-normal modes-driven protein transitions

We predict protein transitions using the NOLB method (68, 82). Starting from a protein 3D structures, NOLB builds an elastic network model where any two atoms closeby in 3D space are linked by a harmonic spring. The model is anisotropic and comprises all the atoms of the protein (101, 102). It has the following potential function,

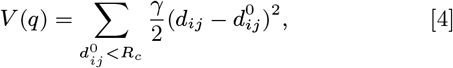

where *d*_*ij*_ is the distance between the *i*^*th*^ and the *j*^*th*^ atoms, 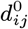 is the reference distance between these atoms, as found in the original structure, *γ* is the spring constant, and *R*_*c*_ is a cutoff distance, set at 5 Å by default. Optionally, NOLB can modify the network by removing the links formed between some pairs of protein regions given as input. We use this functionality to account for the results generated by HOPMA in the construction of the elastic network.

NOLB then diagonalizes the rotation translation blocks (RTB)-projected mass-weighted Hessian matrix of this potential function to compute a set of eigenvectors or normal modes. The RTB technique is used to reduce the dimensionality of the diagonalization problem, by considering individual or several consecutive amino residues are as rigid blocks (103–105). NOLB then deforms the input structure by extrapolating the instantaneous linear velocities and instantaneous angular velocities composing the eigenvectors to a series of twists (82). This guarantees a high stereo-chemical quality of the predicted conformations. To predict a transition from the *input* structure to some *target* structure, NOLB computes the displacement between the two structures and determines the optimal normal mode amplitudes (68). The transition can be optionally decomposed into smaller steps, each corresponding to a total linear deformation RMSD of 0.1 Å. The algorithm stops when the maximum number of iterations is exceeded (100 by default), or if the relative deformation becomes smaller than a tolerance of 1*e*− 6. NOLB also implements an iterative scheme where, at each iteration, a new elastic network model is built, representing the last predicted conformation, and the corresponding normal modes are computed (68).

### Assessing the predicted transitions

To assess the quality of the computed transitions, we measure the extent to which they cover the conformational deviation between the aligned starting and target states. Transition coverage is computed as

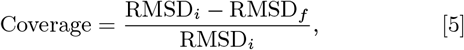

where RMSD_*i*_ is the initial root mean square deviation between the starting and target structures, and RMSD_*f*_ is the deviation between the final structure obtained from the computed transition and the target structure. The coverage varies between 0 (null prediction) and 1 (perfect prediction).

## 3. Results and Discussion

### A colored-map-based approach to alleviate protein structural constraints

We model a protein 3D structure as a graph 𝒢 = (𝒱, *ℰ*), where each node *v* ∈ 𝒱 corresponds to a protein atom and each edge in ℰ connecting *v* to *v*^*′*^ represents an harmonic spring linking the corresponding atoms. The edges’ weights indicate the spring constants. The simplest way to build such a graph, called a protein elastic network, is to determine the set of edges ℰ based on the interatomic pairwise euclidian distances and to set a unique value for the spring constant. Specifically, two nodes *v* and *v*^*′*^ ∈ 𝒱 will be linked by a fixed-weight edge if they are distant by less than a certain distance cutoff, typically comprised between 3.5 Å and 15 Å. Different protein shapes will lead to different network topologies encoding different information about the dynamical potential of the protein.

Here, we propose to build an elastic network that only partially matches the protein structure to maximise its dynamical potential. To decide whether two nodes *v* and *v*^*′*^ should be linked by an edge, we account for the 3D proximity of the corresponding atoms and also for their environment (**Fig. 1A-C**). More precisely, any two connected atoms in the network are closeby in 3D space (distant by less than 5 Å) and located in an extended contact area between two contiguous protein segments. We detect contact areas between protein segments as contiguous patches in a smoothed binarized *contact propensity map* (**Fig. 1B**) computed from the C*α*-C*α* interatomic distances (**Fig. 1A**). These patches can be seen as connected components in a graph encoding 3D proximity at the residue level. In a globular protein, the largest patch is located around the diagonal and corresponds to the contiguous area defined by local contacts along the backbone (**Fig. 1B**, in red). The off-diagonal patches correspond to interactions between disjoint segments remote in the protein sequence (**Fig. 1B**, in other colors). We allow building edges within large patches, representing extended contact areas, and forbid the creation of edges between atom pairs located in small patches or outside of the patches (**Fig. 1C-D**).

**Fig. 1.**
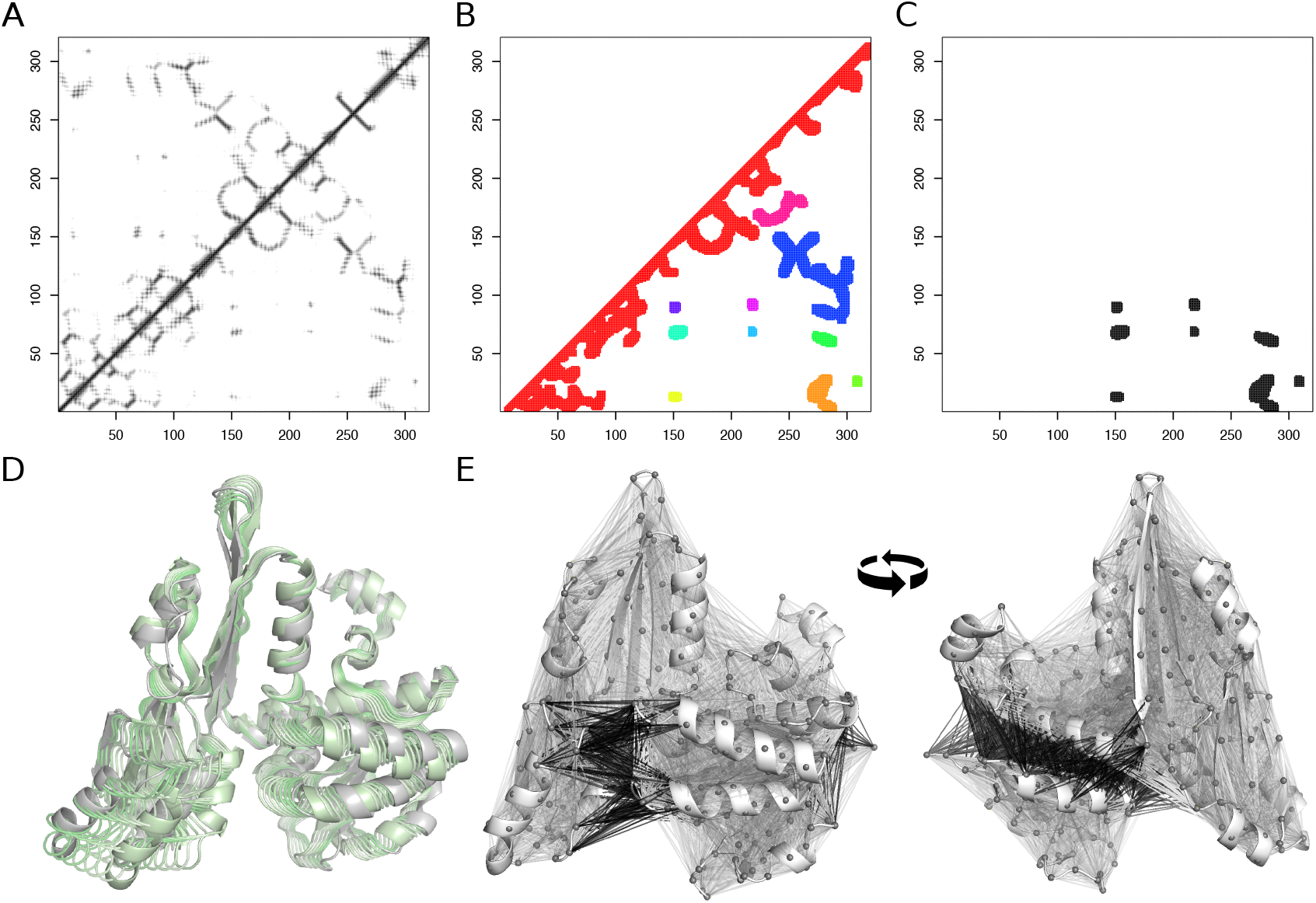
Principle of the method. The closed conformation of the diaminopimelic acid dehydrogenase (PDB code: 1dapB) is taken as an illustrative example. **A**. Distance matrix computed between C*α* atoms. The darker the dot, the closer the residues. **B**. Coloured binarised contact propensity matrix. Each colour indicates a contiguous patch of active cells in the matrix. **C**. Selection of the eight smallest patches. **D**. Nonlinear-normal-mode-driven transition toward the open conformation of the enzyme (PDB code: 3dapA). The initial and intermediate conformations are in green while the target conformation is in grey. The contacts corresponding to the eight identified patches were removed from the initial structure prior to the calculation of the transition. **E**. Mapping of the contact matrix shown in panel B onto the 3D structure of the protein. The contacts corresponding to the eight identified patches are highlighted in black, while the other ones are in grey.

We implemented our approach as an automated tool called HOPMA – proteiN cOloRed MAps. We assessed the ability of HOPMA to reveal protein 3D structures dynamical potential on 402 protein functional transitions covering a large spectrum of motions.

### HOPMA facilitates protein structure opening

As a proof-of-concept, we first applied HOPMA to a set of 23 proteins exhibiting opening-closing motions (**Fig. 2**). We computed the transitions between the closed and open forms, in forward and backward directions, using the very rapid NOnLinear rigid Block (NOLB) NMA. Given a initial-target structure pair, NOLB computed the normal modes of the initial structure and deformed it along a combination of the ten lowest-frequency modes (see *Materials and Methods*). We evaluated the quality of a predicted transition by computing its transition coverage as the percentage of RMS deviation between the initial and target structures covered by the prediction.

**Fig. 2.**
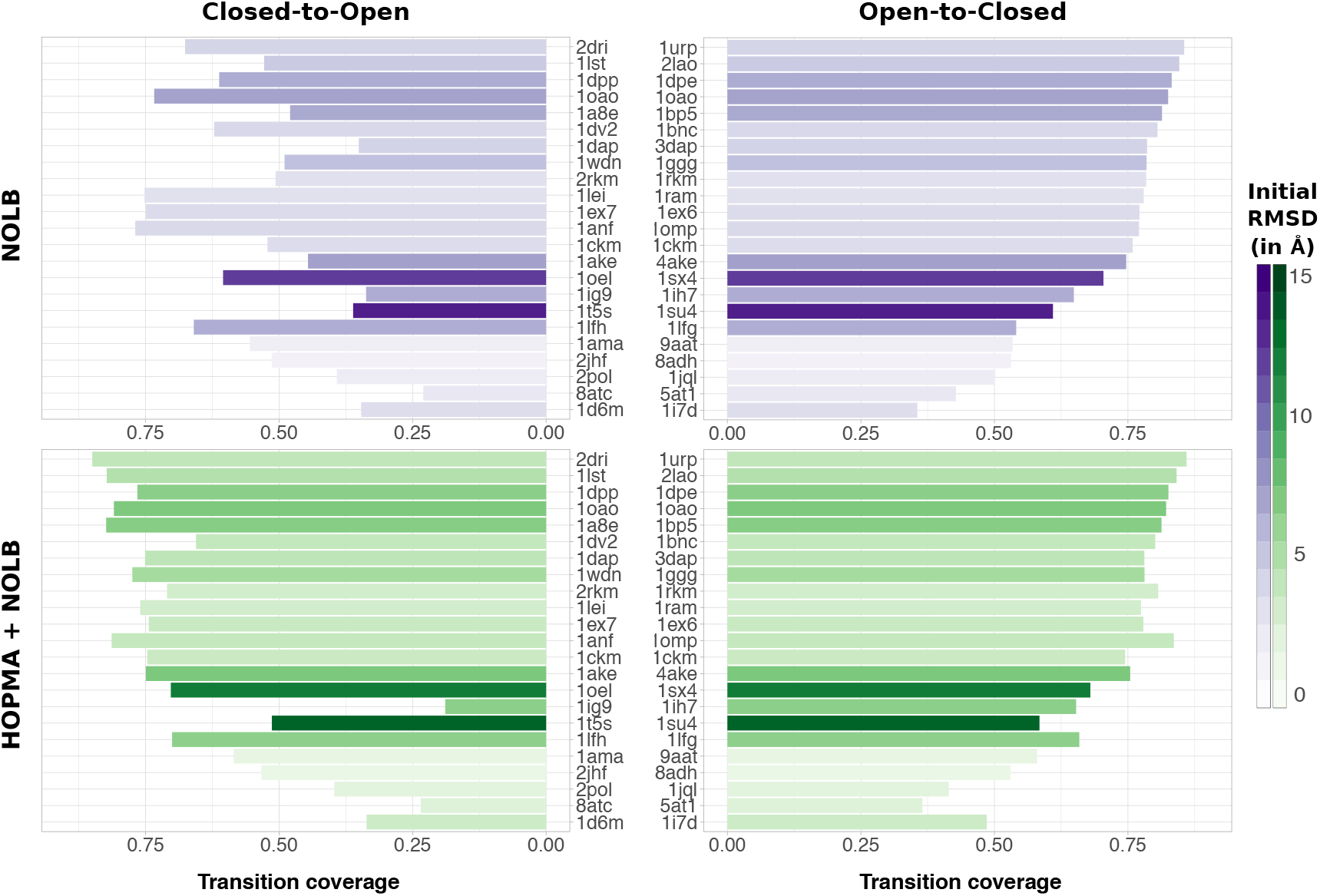
Transition coverage for opening/closing motions. Comparison of the coverages achieved by the NOLB nonlinear modes for 23 transitions (see *Materials and Methods*, iMod benchmark) between a closed state and an open state, in the forward direction (closed-to-open, on the left) and backward direction (open-to-closed, on the right). On top, all the atom pairs distant by less than 5 Å are linked by a spring. At the bottom, we used HOPMA to remove some links. The darker the color, the higher the RMSD between the open and closed structures.

Figure 2 shows a double comparison, the first one between the closed-to-open (on the left) and open-to-closed (on the right) directions, the second one between the “classical” elastic network (in purple), where the links are defined using a distance cutoff (5 Å), and the modified network (in green), where some links are removed according to HOPMA results. Accounting for HOPMA results in the definition of the elastic network (**Fig. 2**, in green) allows achieving a transition coverage in the closed-to-open direction almost equivalent to that of the open-to-closed direction (average Δ*Coverage* = 5 *±* 10%). By contrast, the coverage difference between the two directions is of 16% on average when starting from the “classical” elastic networks (**Fig. 2**, in purple). The HOPMA-enhanced networks are substantially easier to open, leading to an average coverage of the transition to the open form of 65 *±* 19%, compared to 53 *±* 15% with the “classical” elastic networks (**Fig. 2**, *Closed-to-Open*). The highest improvement is observed for the diaminopimelic acid dehydrogenase (1dap-3dap) for which the coverage increases from 35% to 75% (**Fig. 1**). Moreover, HOPMA leads to a significant coverage improvement (by more than 5%) in the open-to-closed direction for three proteins (**Fig. 2**, *Open-to-Closed*), namely the maltodextrin-binding protein (1anf-1omp), the dna topoisomerase III (1d6m-1i7d), and the lactoferrin (1lfh-1lfg).

### HOPMA improves the prediction of a wide range of motions

We then assessed HOPMA on a large and diverse testing set (AlterBench) comprising 143 protein structure pairs with RMS deviations ranging from 3 to 35 Å (**Fig. 3A**). All transitions were computed in the forward and backward directions, except for four cases, leading to a total of 282 transitions (see *Materials and Methods*). For the vast majority (98%) of cases, the HOPMA-enhanced networks lead to equivalent or better results compared to the “classical” networks (**Fig. 3A**). Moreover, HOPMA helps to describe a wide range of motions, including opening/closing motions, motions involving domain rotations, concerted motions, and some more complex motions.

**Fig. 3.**
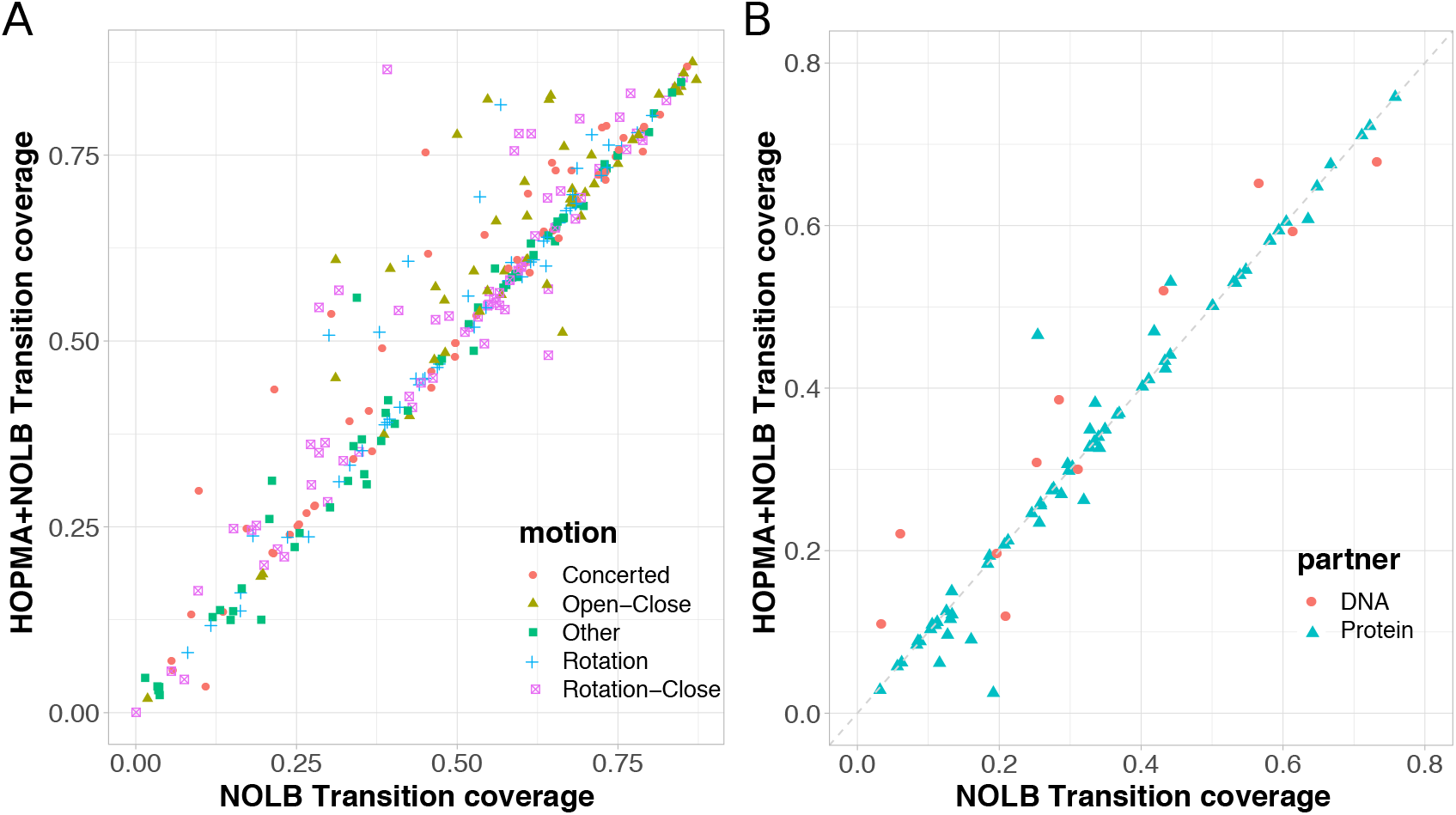
Large scale assessment of HOPMA. The transition coverages achieved with the elastic networks defined using HOPMA are compared with those obtained with the “classical” networks (distance cutoff only). **A**. 282 transitions displaying a wide range of motions from the AlterBench dataset (see *Materials and Methods*). For each pair of structures defined in the set, we computed both the forward and backward transitions. **B**. 79 unbound-to-bound transitions from the PPDBv5 (protein-protein interactions) and HR-PDNA187 (protein-DNA interactions).

For example, HOPMA substantially improves the prediction of a rotation-close motion occurring in the pilus retraction motor PilT (2gsz), in both the forward and backward directions (**Fig. 4**). This hexameric ATPase is required for bacterial type IV pilus retraction and surface motility. In the hexamer, the ATPase core of one subunit extensively interacts with the N-terminal domain of the next subunit. As a result, the inter-domain distance and orientation vary from one subunit to another. The modelled transition takes place between two subunits (chains B and C) and implies some rearrangements of the contacts between the two domains (**Fig. 4**, compare the two colored maps). HOPMA identifies and removes the contacts between the two domains in both structures, leading to an increased transition coverage, from 39% to 86% in the forward direction (**Fig. 4**, on top), and from 62% to 78% in the backward direction (**Fig. 4**, at the bottom).

**Fig. 4.**
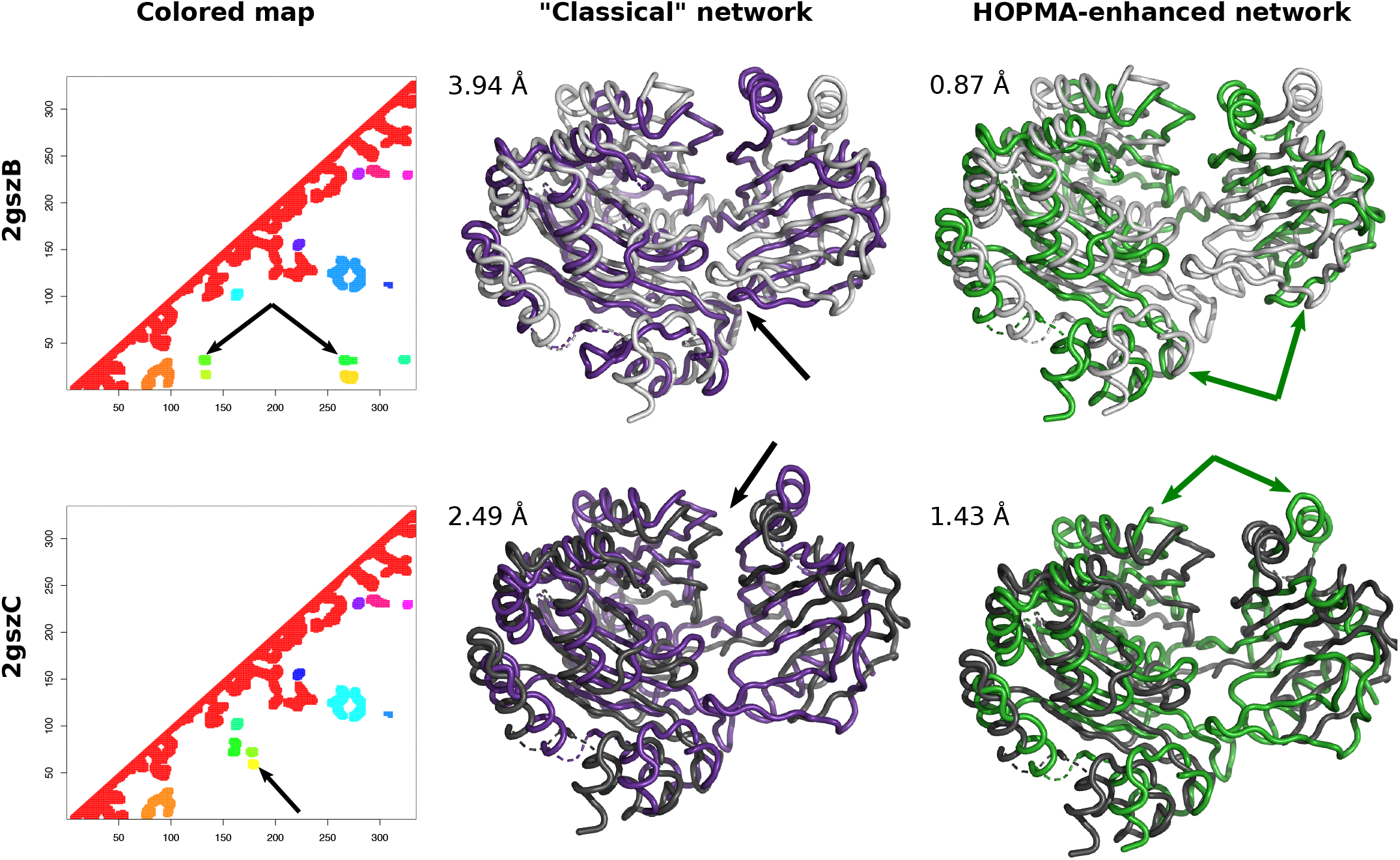
Conformational change between two subunits of the pilus retraction motor PilT. The experimental structures of the two subunits are shown as light grey (2gszB, on top) and dark grey (2gszC, at the bottom) cartoons. They deviate by 6.47 Å. The corresponding coloured maps computed by HOPMA are displayed on the left. The final conformations of the predicted transitions are shown as purple (“classical” network) and green (HOPMA-enhanced network) cartoons and superimposed onto the initial structure. Their RMS deviations to the target structures are given. The black arrows indicate some contacting regions in the initial structures, which are maintained in the conformations predicted from the “classical” networks. The green arrows highlight some opening motions correctly predicted in these regions when starting from the HOPMA-enhanced networks.

We also tested HOPMA on two other datasets, namely PPDBv5 and HR-PDNA187, dedicated to the study of protein-protein and protein-DNA interactions respectively (**Fig. 3B**). We selected 79 unbound-bound structure pairs with at least 2 Å RMS deviation between them, and we only focused on the prediction of the unbound-to-bound transitions (see *Materials and Methods*). As observed on the AlterBench set, the HOPMA-enhanced elastic networks lead to equivalent or better transition coverages compared to the “classical” elastic networks (**Fig. 3B**). However, the transitions are overall more difficult to predict compared to the AlterBench set (**Figure S1**). HOPMA significantly improved the predictions of nine functional transitions, three undergone upon binding to a protein partner and six upon binding to DNA (**Fig. 3B**). For example, it increases the coverage of the transition of the human vinculin head upon binding to the vinculin tail from 25% to 47%. The conformational change between the initial and target structures concern 20% of the protein and amounts to 4.25 Å. The low coverage obtained with the “classical” elastic network can be explained by the presence of contacts with Cterminal extremity the locks the position of a helix-turn-helix. In the HOPMA-enhanced network, the contacts are removed and the helices are free to rotate (**Fig. S2**).

### HOPMA reduces the number of iterations required for very large DNA-induced conformational changes

To describe very large transitions or transitions involving complex motions, it may be useful to re-iterate the NOLB calculation one or several times. At each iteration, a new elastic network model is computed from the last predicted conformation and a new transition is computed using its first ten normal modes (see *Materials and Methods*). We investigated whether HOPMA could also be useful in this multi-iterative scheme. We focused on the 11 transitions associated with DNA binding from the HR-PDNA187 dataset. We found that accounting for HOPMA results to define the elastic networks leads to conformations that are closer to the target, after three iterations, for the three very large transitions of the set (**Fig. S3**, initial RMSD>10 Å). The highest gain is obtained for the Y-family DNA polymerase Dpo4 (2rdiA-1jxoA). This enzyme plays a crucial role in translesion synthesis and undergoes a dramatic conformational change upon DNA binding (**Fig. 5**). The transition implies a ∼ 130 degrees rotation of the little finger domain relative to the polymerase core resulting in a RMS deviation of 11.47 Å between the unbound and bound structures. HOPMA marked the contacts between the two domains as to be removed and this resulted in a transition coverage of 38% instead of 28% with the “classical” network after the first iteration. Re-iterating the transition computation twice produces a final conformation 1.80 Å away from the target that can accommodate DNA without backbone clashes (**Fig. 5**, in purple). By contrast, the same procedure without using HOPMA produces a more distant conformation, with visible distortions in the little finger domain, that cannot accommodate DNA (**Fig. 5**).

**Fig. 5.**
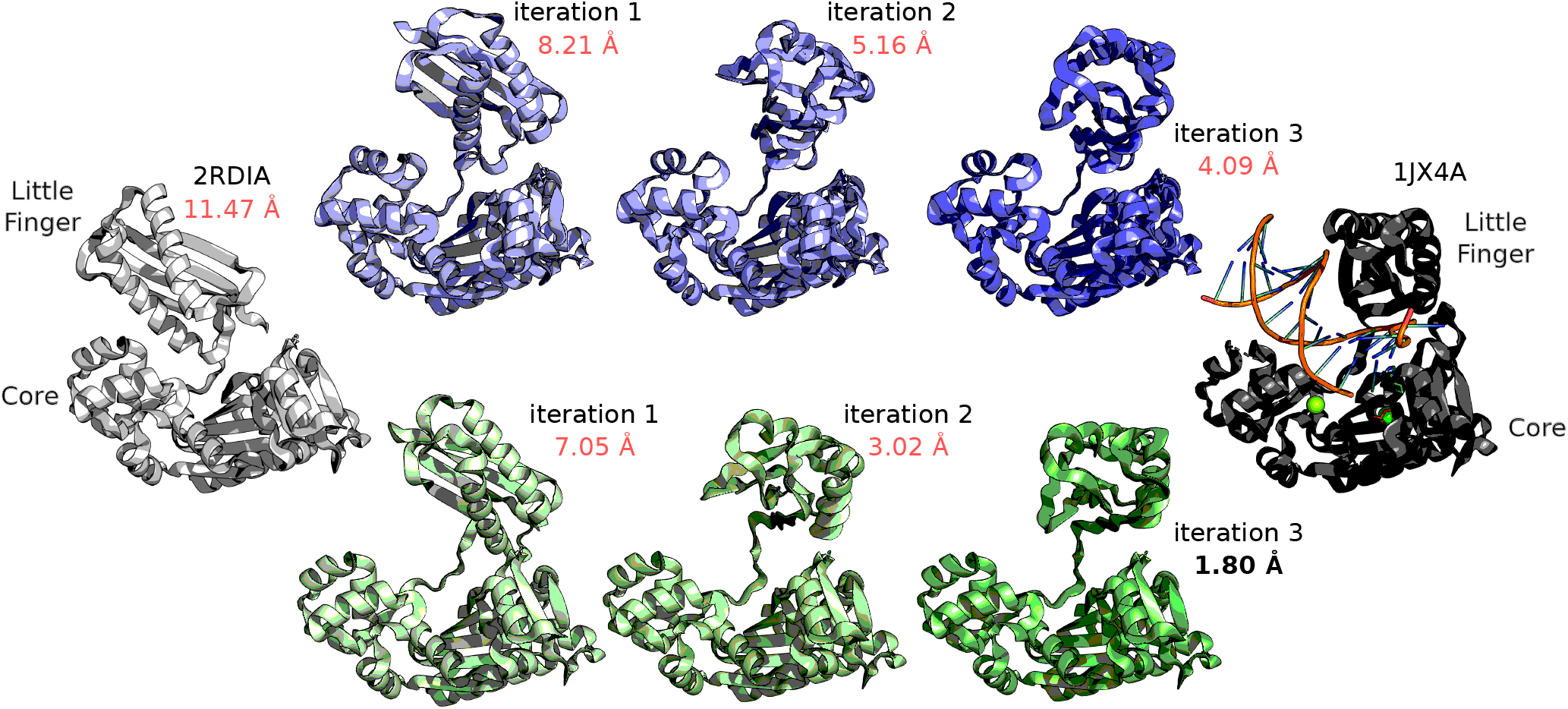
DNA Polymerase conformational change upon binding to DNA and an incoming nucleotide. The unbound structure (2rdiA) is shown on the left in grey while the structure bound to DNA and a nucleotide (1jx4A) is shown on the right in black. The little finger and core domains are labeled. The conformations in between were predicted by NOLB, after 1, 2 and 3 iterations respectively. The conformations in purple were obtained from “classical” elastic networks, while those in green were obtained from less constrained networks, where the links identified by HOPMA were removed. The RMS deviations to the target are given and highlighted in red when the backbone of the conformation clashes with the DNA.

### HOPMA can be useful to explore a protein conformational space

Beyond the prediction of a transition between two functional states, we investigated whether HOPMA could help to navigate within an ensemble of protein conformations. We took the multi-functional calcium sensor calmodulin as a case study. Calmodulin structural flexibility has been extensively characterized and shown to play a key role in its ability to bind a tremendous amount of proteins and peptides (106). We computed all the pairwise transitions between 27 conformations of the protein, both in the forward and in the backward directions (**Fig. 6**). The HOPMA-enhanced networks systematically led to equivalent or better predictions compared to the “classical” networks (**Fig. 6B**). One can clearly identify seven conformers (numbered 1, 5, 12, 18, 20, 25 and 27) whose transitions to the other conformers are significantly facilitated. In these conformers, the two domains of the protein establish substantial contacts that prevent them from changing their position and orientation with respect to each other. HOPMA detects the contacts and marks them as to be removed, enhancing the dynamical potential of the protein network. The most striking coverage gain is observed for the conformer 27, which corresponds to an X-ray crystallographic structure of the protein complexed with a substrate peptide and four calcium ions (**Fig. 6C-D**). The conformation is very compact, with three contact areas between the two protein domains (**Fig. 6C**). While NOLB is unable to predict the transitions to the other conformers when starting with the “classical” networks (average *Coverage* = 0.22), it covers 50% of the transitions, on average, when starting from the less constrained HOPMA-enhanced network (**Fig. 6D**, see also **Fig. S4**). Nevertheless, this is still significantly less that what is achieved among the others conformers, which correspond to free forms of the protein (average *Coverage* = 0.73, see **Fig. S4**). This difference may be explained by more subtle and more difficult to predict conformational changes between the free and complexed states that affect the orientation of the helix pairs (EF-hand motifs) within each domain and the associated solvent exposure of some hydrophobic residues.

**Fig. 6.**
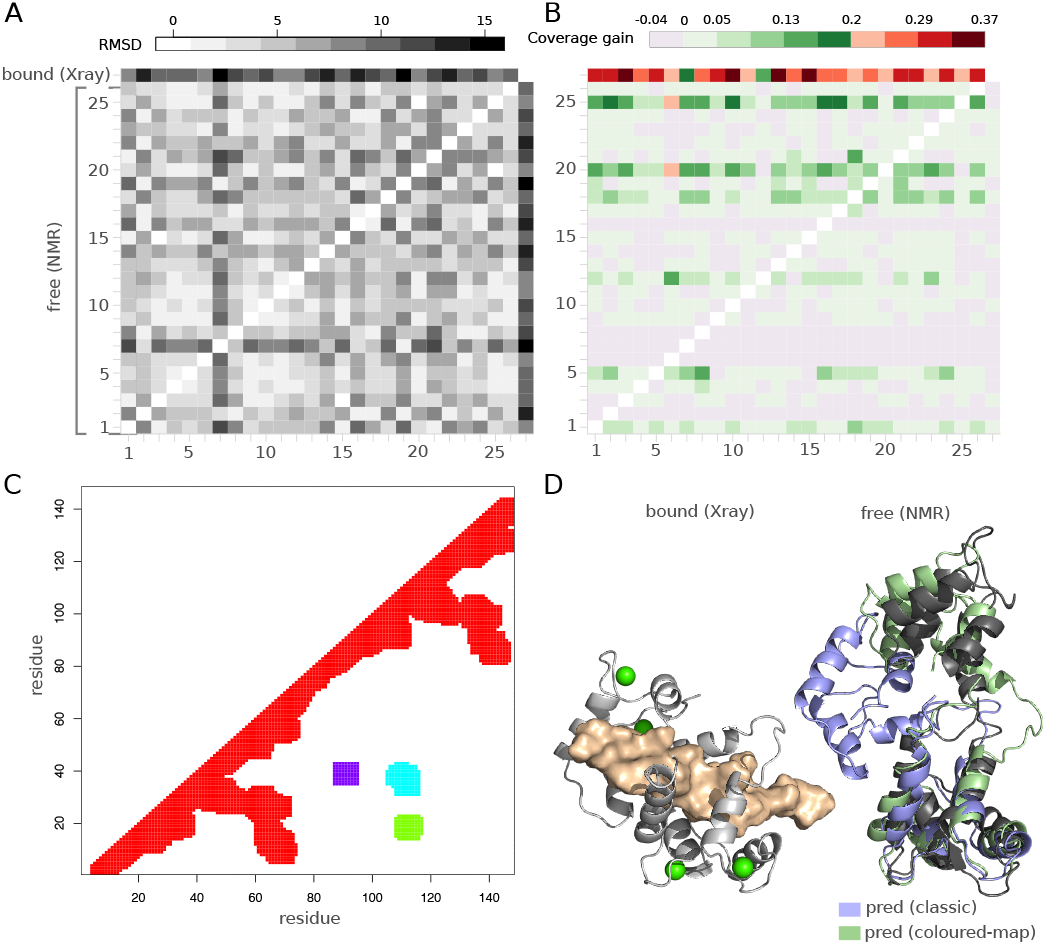
Exploration of Calmodulin conformational space. **A**. Pairwise RMS deviations between 27 conformations of calmodulin, corresponding to 26 NMR models of the free protein (1cfc and 1cfd) and one crystallographic structure of the protein bound to a peptide and some calcium ions (2bbm). **B**. Transition coverage gain when predicting all pairwise transitions starting from HOPMA-enhanced elastic networks with respect to “classical” elastic networks. **C**. Colored map computed by HOPMA for the bound structure (2bbm). **D**. On the left, the bound structure is displayed as a light grey cartoon with the peptide in colored surface and the calcium ions as green spheres. On the right, the free structure (first monomer of 1cfc) is displayed as a dark grey cartoon, while the final conformations predicted from elastic networks representing the bound structure are displayed as colored cartoons. The conformation predicted from the “classical” network is shown in purple while the one generated from the HOPMA-enhanced network is in green.

### Computational details and availability

The HOPMA code was written in Python3 and R. All the calculations were performed on a 2,7 GHz Intel Core i7 CPU with 16 Go RAM. Given a protein structure, we used the following command to run HOPMA: “python3 $HOPMA_PATH/hopma.py *prot* -c *chain*”, where *prot* is the name of the input PDB structure file without the extension, and *chain* is the name of the chain to analyze. Given two protein structures *protInitial* and *protTarget*, we used the following command to run NOLB: “NOLB *protInitial protTarget* -n 10 --nlin -m”. To account for HOPMA results, we used the “--excl” option with the output of HOPMA. The computations are efficient, for example, predicting a transition for a protein of about 300 residues takes 12 seconds, including 9 seconds to perform HOPMA analysis and 3 seconds to run NOLB. The execution time increases up to 81 seconds for a protein of about 1 000 residues (66 seconds for HOPMA and 15 seconds for NOLB). The efficiency of HOPMA can be improved by transferring it to C++. The method is available at https://github.com/elolaine/HOPMA and https://team.inria.fr/nano-d/software/nolb-normal-modes.

## 4. Conclusions

This work further develops our approach to predicting conformational changes in proteins and other macromolecules. Here, we implemented and carefully assessed the novel idea of modifying the initial elastic network by removing links corresponding to disconnected patches in the contact map. To detect these patches, we rely only on the 3D information encoded in the input protein structure. As expected, the modified networks led to significantly improved transitions, as we showed on a large set of examples taken from four diversified benchmarks. The method is user-friendly and computationally efficient. This work opens perspectives for sampling new protein functional conformations that were not accessible before, starting from experimentally determined structures.

## 5. Acknowledgments

The authors are thankful to Vladimir Sorokin for inspiration.

