## Supplementary Figures S1-S4 for "HOPMA: Boosting protein functional dynamics with colored contact maps"

### Supporting Information for HOPMA: Boosting protein functional dynamics with colored contact maps

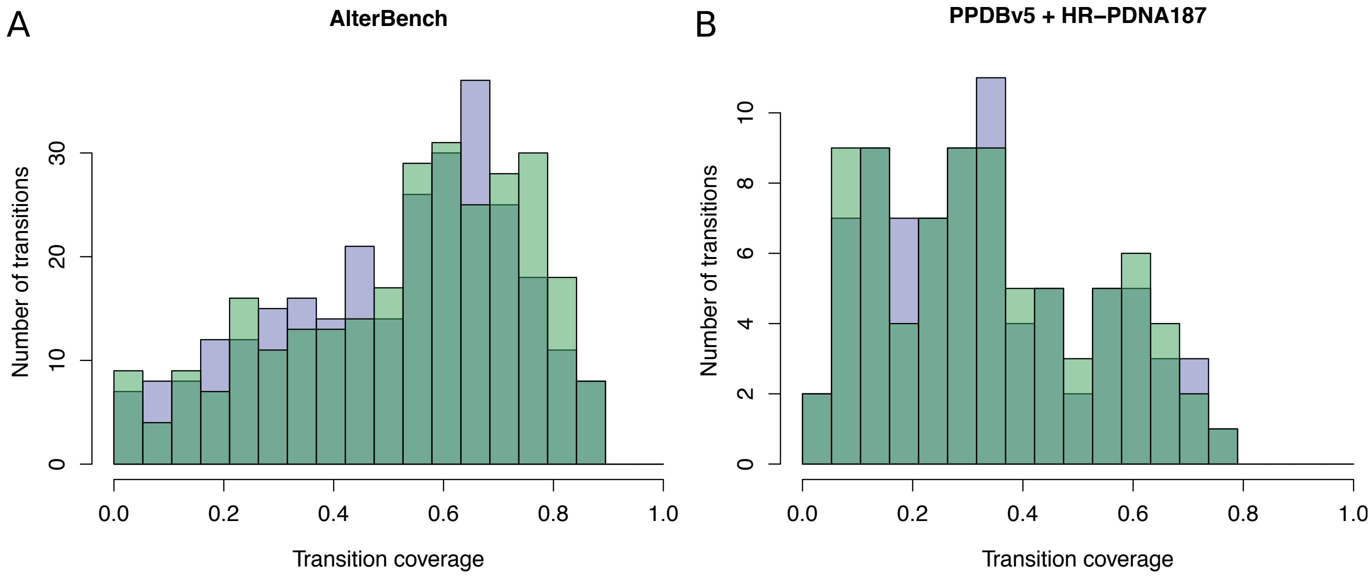


Figure S1: **Transition coverage distributions.** The distributions in purple correspond to the coverages obtained when starting from the "classical" elastic networks, while the green distributions give the coverages obtained from the HOPMA-enhanced networks. **A.** 282 transitions displaying a wide range of motions from the AlterBench dataset (see *Materials and Methods*). For each pair of structures defined in the set, we computed both the forward and backward transitions. **B.** 79 unbound-to-bound transitions from the PPDBv5 (protein-protein interactions) and HR-PDNA187 (protein-DNA interactions).

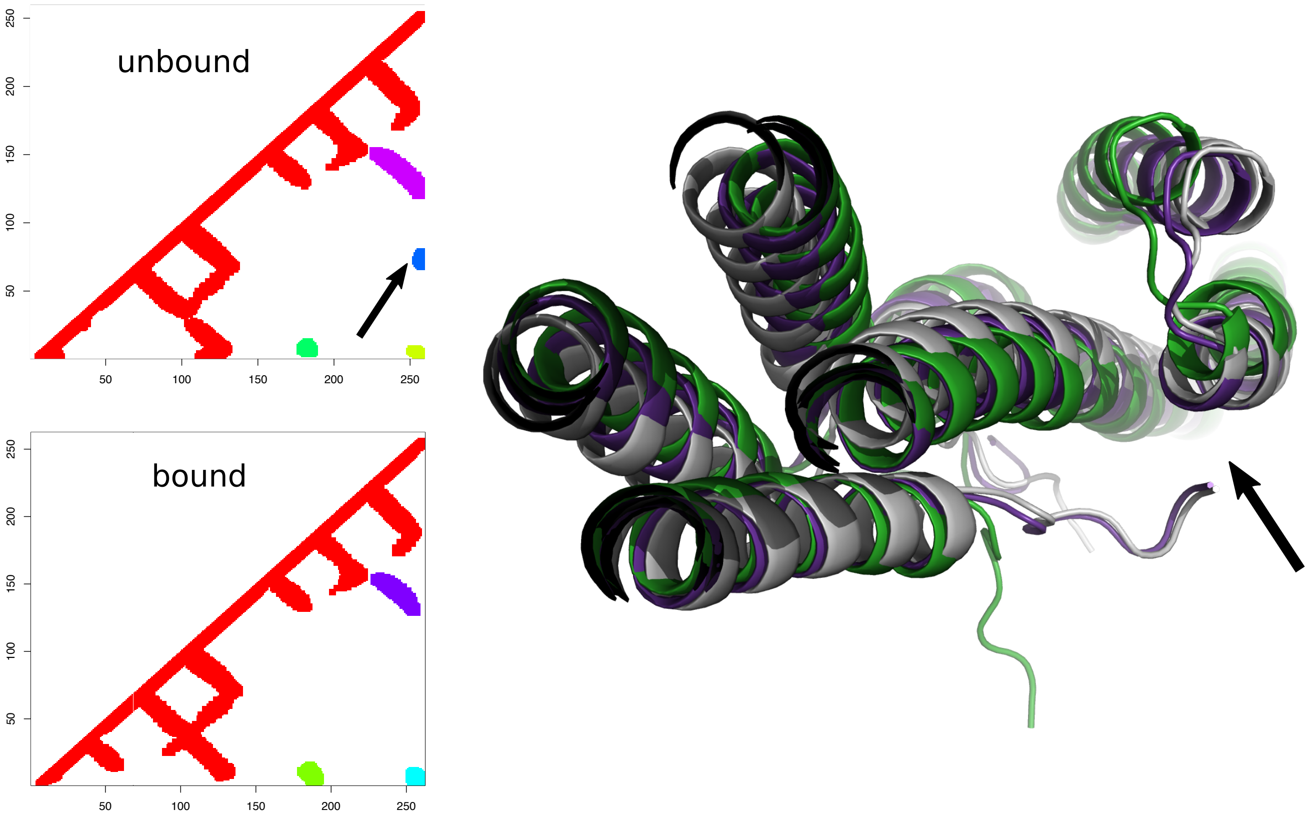


Figure S2: **Transition of the the human vinculin head upon binding to the vinculin tail.** The colored maps computed for the unbound and bound conformations are shown on the left. The protein structure is shown on the right as a cartoon. The unbound conformation is displayed in light gray. The conformations predicted by NOLB are show in purple and green. The purple one was obtained from the "classical" elastic network representing the unbound conformation, while the green one was obtained from the HOPMA-enhanced network. The arrows indicate a contact area between the C-terminal extremity and a helix undergoing a rotation between the two conformations.

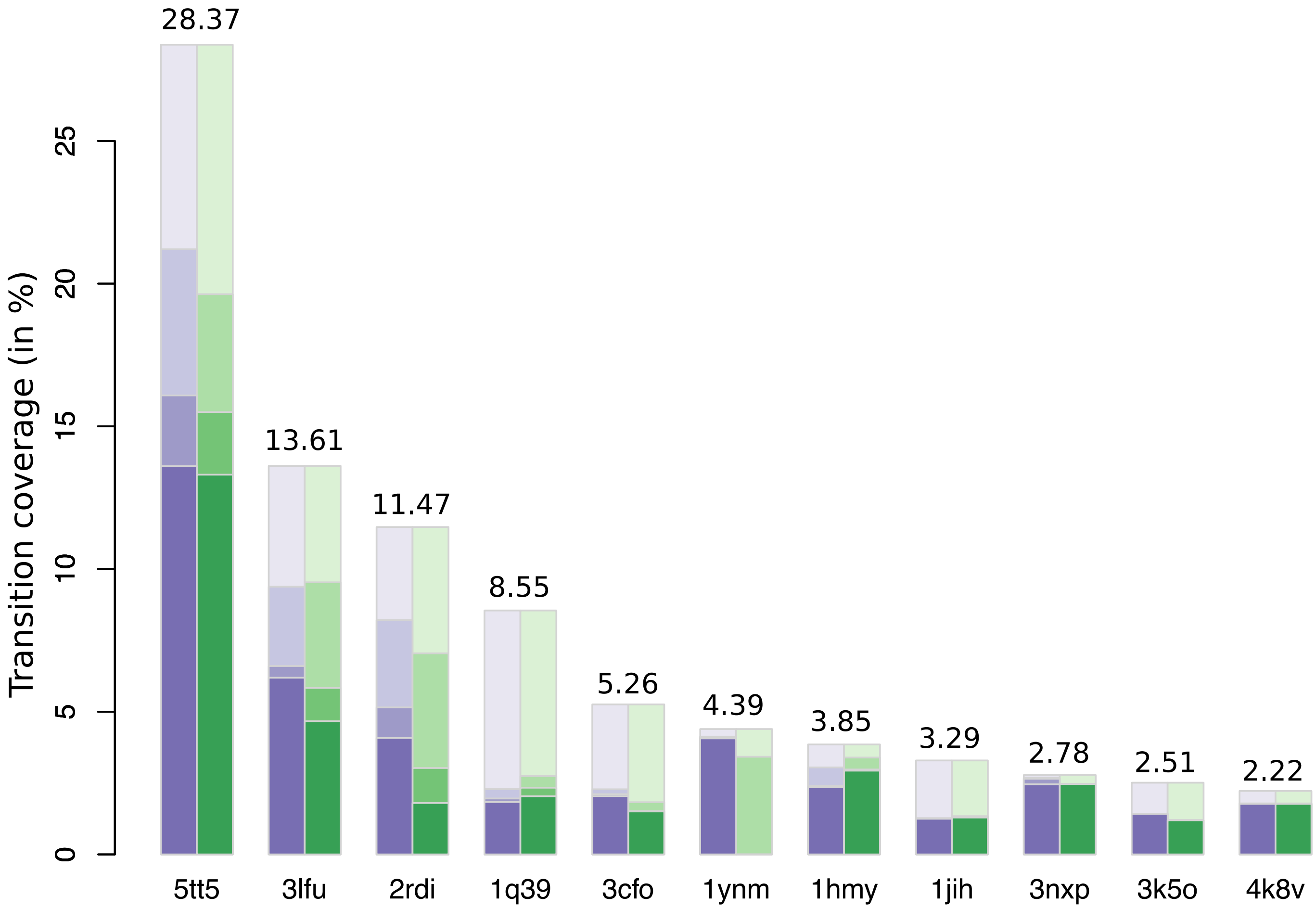


Figure S3: **Transition coverages achieved with 3 iterations for the DNA-induced conformational changes.** The PDB codes of the source structures are indicated in x-axis. The RMSDs to the target are indicated on top of the bars, in Å. Each bar reports the coverages achieved after 1, 2 and 3 iterations of NOLB (in purple) or of HOPMA+NOLB (in green) in different color tones.

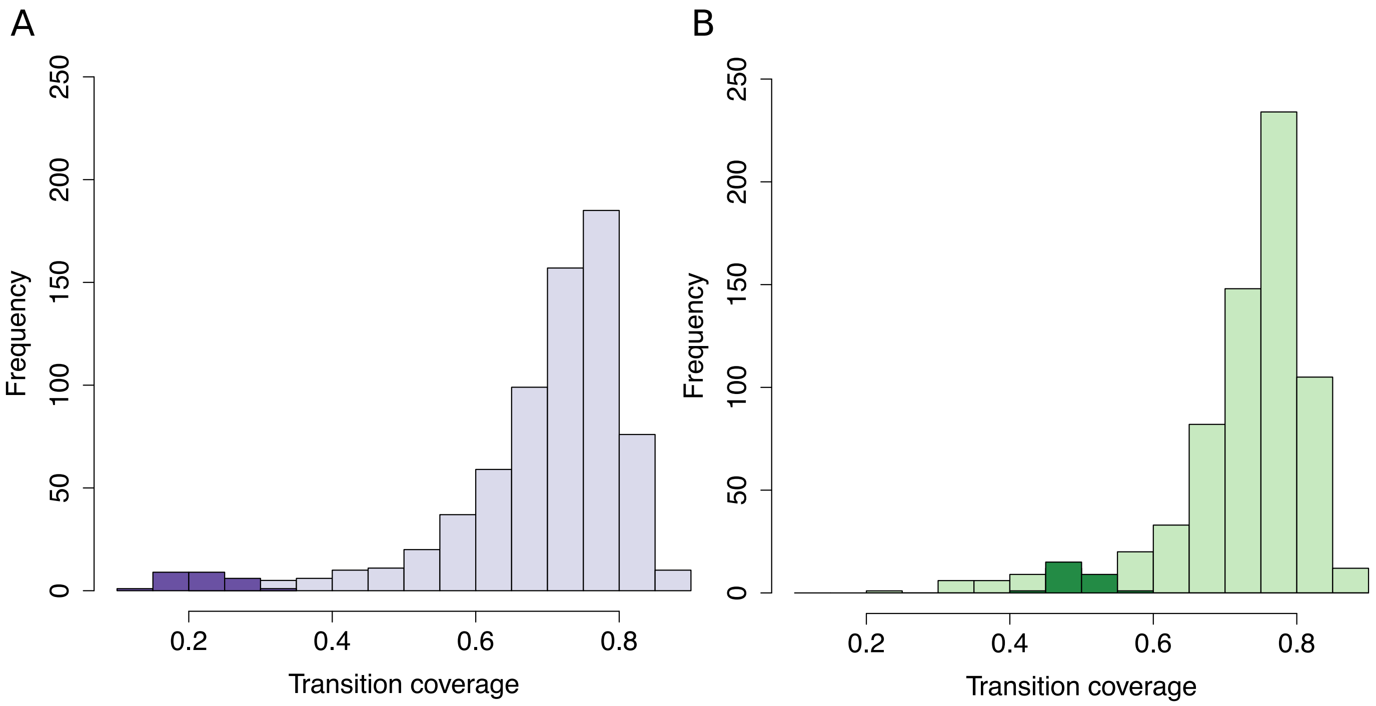


Figure S4: **Transition coverages for calmodulin conformational ensemble.** Each distribution comprises 702 values, corresponding to all transitions, in both forward and backward directions, between 27 conformers of the calmodulin. The distribution in purple was obtained from the "classical" elastic networks (A), while the one in green was obtained from the HOPMA-enhanced networks (B). The parts highlighted in dark colors correspond to the transitions from the peptide-bound compact X-ray structure (2bbm) to the other conformations, all determined by NMR (1cfc and 1cfd).
